# An equilibrium binding model for CpG DNA-dependent dimerization of Toll-like receptor 9

**DOI:** 10.1101/573238

**Authors:** Stephanie Reikine, Stephen H. McLaughlin, Christopher M. Johnson, Yorgo Modis

**Author notes:** Correspondence: Yorgo Modis.

## Abstract

Microbial nucleic acids in the extracellular milieu are recognized in vertebrates by Toll-like receptors (TLRs), one of the most important families of innate immune receptors. TLR9 recognizes single-stranded unmethylated CpG DNA in endosomes. DNA binding induces dimerization of TLR9 and activation of a potent inflammatory response. To provide insights on how DNA ligands induce TLR9 dimerization, we developed a detailed theoretical equilibrium ligand binding model. Light scattering and fluorescence polarization assays performed with a recombinant TLR9 ectodomain fragment and a panel of agonistic and antagonistic DNA ligands provide data that restrain the binding parameters in our binding model. This work brings us one step closer to establishing a rigorous biochemical understanding of how TLRs are activated by their ligands.

## Introduction

The first line of defense that vertebrates rely on when presented with a pathogen is the innate immune system [1]. Innate immune receptors have evolved to detect pathogen-associated molecular patterns (PAMPs), chemical motifs that are common in microbes, absent in the host and cannot be readily mutated. A major family of innate immune receptors is the Toll-Like Receptors (TLRs) [2]. TLR3, TLR7, TLR8, and TLR9 are all found in endosomes and recognize nucleic acid PAMPs [3-7]. TLR9 recognizes single-stranded DNA (ssDNA) oligonucleotides containing an unmethylated CG nucleotide sequence motif (CpG) [7]. This type of genomic signature is more prevalent in bacteria and viruses than it is in the mammalian genome, where the majority of CG sites are methylated [8, 9].

The crystal structures of TLR9 ectodomain fragments from mouse, horse and cow have been determined without ligand (apo), bound to antagonistic ligands 4084 and iSUPER, and bound to a truncated version of the activating oligonucleotide ligand 1668 (described previously [10]), termed 1668-12mer [11]. These structures provided the structural basis for the CpG specificity of TLR9 ligand recognition. The CpG motif is directly recognized by several TLR9 amino acids in the binding groove, forming hydrogen bonds and water-mediated interactions [11]. The TLR9 ectodomain crystallized as a monomer without ligand but formed a dimer with 1668-12mer bound, suggesting a model of TLR9 signal activation through dimerization. In the context of the full-length membrane-inserted receptor TLR9 is thought to form loosely assembled inactive homodimers prior to binding ssDNA, with ligand binding inducing a conformational rearrangement and tightening of the dimer assembly necessary to activate signaling [12]. The two TLR9 ectodomains in the agonist-bound structure assemble around two 1668-12mer oligonucleotides to form a 2:2 TLR9:oligonucleotide complex [11]. The oligonucleotides, sandwiched between the two ectodomains, function as ‘molecular glue’ between the two TLR9 subunits [11]. Each ligand in the dimer interacts with two distinct binding surface on TLR9, near the N-and C-terminal ends of the ectodomain, respectively [11]. An additional binding site in TLR9 was recently identified in the central region of the ectodomain, with specificity for short ssDNA accessory oligonucleotides containing the motif 5’-xCx [13], which function as auxiliary ligands to enhance signaling [14]. Auxiliary ligands with analogous functions in signal augmentation have been identified for TLR7 [15] and TLR8 [16].

Although structural studies have shed light on how TLR9 recognizes ssDNA ligands, key open questions remain concerning the signaling mechanism of TLR9. A reductionist approach to determine the minimal sequence requirements for an oligonucleotide to maximally activate TLR9 identified a length of between 23 and 29 nucleotides as the optimal length for mouse TLR9 agonists, including a 5’-TCC motif and CpG motif located 5-7 nucleotides from the 5’ end [17, 18]. It remains unclear why extending the length of the ligand beyond the 12 nucleotides observed in the TLR9:1668-12mer structure enhances signaling. Moreover, modelling studies of ssDNA have been limited by the use of either a 1:1 binding model (rather than a 2:2 model) or of the Hill equation [11, 13, 19, 20]. Although it is known that the final state of a signaling competent TLR9-ssDNA complex is a 2:2 dimer, it is not known through which pathway this dimerization occurs. TLR9 dimerization upon ligand binding can theoretically occur via two pathways: (1) two TLR9 ectodomains first form 1:1 protein:ligand complexes, which then come together to form 2:2 dimers, or (2) a single TLR9 first binds two oligonucleotides (one at each binding site), and this 2:1 complex then recruits a free TLR9. Determining the primary pathway of TLR9 dimerization and measuring binding cooperativity would provide key missing links in our understanding of TLR9 activation. Here, we propose an equilibrium binding model for ligand-dependent dimerization of TLR9. We support and refine our model with biochemical and biophysical analyses of ligand binding. Our work brings us one step closer to establishing a detailed and rigorous understanding of the kinetic pathway of DNA-dependent TLR9 activation. Given the structural and mechanistic similarities to other TLRs, most notably TLR7 and TLR8, this work could also serve as a more generic model for TLR activation.

## Results

### An equilibrium ligand binding model for TLR9

To generate a complete and quantitative description of ligand-induced dimerization of TLR9, we first need to establish a model of the pathway to dimerization that can be tested experimentally. A stoichiometric binding equilibrium model representing the possible pathways to ligand-induced TLR9 dimerization is presented in **Fig. 1A**. The model allows for assembly of the 2:2 active TLR9:DNA complex via initial dimerization of TLR9 upon binding two ligands, a single ligand or no ligands (**Fig. 1A**). It is important to note that the term [PD] represents the apparent binding of a DNA oligonucleotide (D) to TLR9 (P). Since TLR9 has two oligonucleotide binding sites, [PD] is the sum of [P_A_D] and [P_B_D], representing the oligonucleotide bound to each of the two binding sites, P_A_ and P_B_. Hence, the macroscopic equilibrium constants K_1_, K_2_ and K_3_ are each comprised of at least two microscopic binding constants, which describe the equilibria between [P_A_D] and [P_B_D] and the previous or subsequent state. These macroscopic binding constants also include other microscopic constants if binding induces a conformational change or is cooperative.

**Fig. 1.**
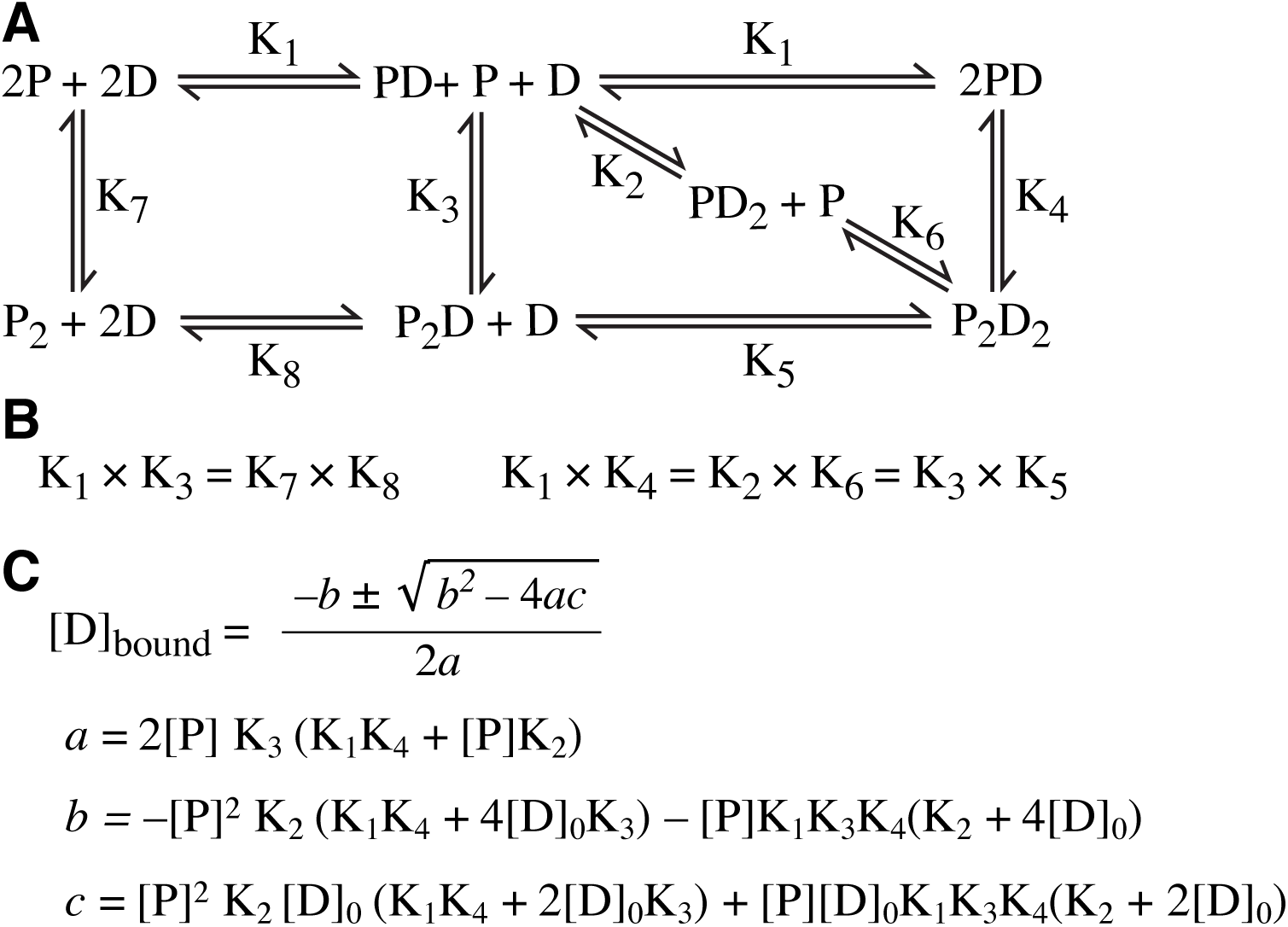
Proposed general equilibrium model for TLR9 agonist binding. (**A**) A stoichiometric representation of the possible species in equilibrium as TLR9 binds an activating ssDNA ligand and dimerizes. The macroscopic equilibrium binding constants are labeled. (**B**) Relationships between the macroscopic constants. (**C**) Solution for the concentration of bound ssDNA, [D]_bound_.

Our model has implications for the relationships between the macroscopic binding constants that are informative (**Fig. 1B**). Since mTLR9 ectodomain (mTLR9-ECD) remains predominantly monomeric even at high protein concentrations [11], our model predicts that K_7_, K_8_ or both are large relative to K_1_. Moreover, given that K_1_K_3_ = K_7_K_8_, K_3_ must be very large. Similarly, since final product is known to be a 2:2 dimer, K_1_K_4_ must be relatively small, which implies that K_5_ must be very small, since K_1_K_4_ = K_3_K_5_ and K_3_ is very large. This leads to the interesting conclusion that the 2:1 TLR9:DNA dimer species (P_2_D) rarely occurs. Our general model of equilibrium binding can be solved for the concentration of bound ssDNA, [D]_bound_, accounting for mass action (**Fig. 1C**). [D]_bound_ be measured experimentally in a ligand binding assay. A complete solution of all macroscopic constants is not readily accessible experimentally, but numerical solutions or simulations could in principle be used to identify possible values for each constant.

Although our equilibrium binding model is expressed in terms of macroscopic binding constants, considering its implications for the microscopic constants is informative. First, we considered the scenario where the two ssDNA binding sites are independent and not cooperative. The microscopic binding constants for K_1_ are K_A_ and K_B_, describing DNA binding to sites P_A_ and P_B_, respectively. Writing K_1_ in terms of the microscopic constants:

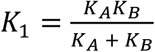

For K_2_, the microscopic binding constants are also K_A_ and K_B_, provided that ligand binding at one site does not alter the binding affinity at the second site, for example through a conformational change in TLR9 or other allosteric mechanism. Writing K_2_ in terms of the microscopic constants, K_2_ = K_A_ + K_B_. The macroscopic constants K_1_ and K_2_ are therefore related as follows:

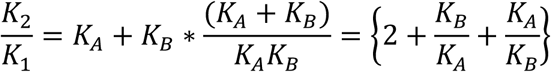

Since the constants cannot be negative, it follows that K_2_ > K_1_. This analysis suggests that the most likely pathway for the ligand-induced assembly of TLR9 dimers is first through saturation of one of the binding sites with ligand to form 1:1 protein:DNA (PD) complexes, which then assemble into 2:2 dimers (P_2_D_2_). However, this analysis assumes that the two binding sites are independent. If this is incorrect and binding of the second ligand is cooperative, K_2_ could be smaller than K_1_.

The microscopic binding constants for K_3_ are more complex than for K_1_ and K_2_. The binding affinity of a free protein to a DNA that is part of a protein:DNA complex is different than its binding affinity to free DNA. Additionally, protein:protein interactions may promote the K_3_ transition. In summary, our theoretical analysis of TLR9 ligand binding based on a specific set of assumptions makes testable predictions, for example K_3_ > K_1_ and K_2_ > K_1_, and provides a framework for experimental characterization of ligand-induced TLR9 dimerization.

### Agonistic ssDNA ligands induce TLR9 dimerization

Having established an equilibrium binding model, we set out to test it and measure key parameters experimentally with recombinant mTLR9-ECD and selected DNA ligands. Oligonucleotides 1668 and 1668-12mer were shown previously to induce dimerization of mTLR9 [11]. Other agonistic ligands are thought to activate TLR9 in the same manner [21], but a systematic comparison of the effect of different ligand on the oligomeric state of TLR9 has not been performed. To address this, we measured the oligomeric state of recombinant mTLR9-ECD in the presence of five different oligonucleotides (**Table 1**) by dynamic light scattering (DLS). Our panel of ligands included the prototypical activating oligonucleotides 1668 and 2006 [10, 22]; the 1668-12mer oligonucleotide used in the structural studies [11]; minM, an oligonucleotide identified in cell-based assays as the minimal sequence required for potent activation of mouse TLR9 [17, 18]; and antagonistic oligonucleotide 4084, as a control for binding without dimerization [11].

**Table 1.**
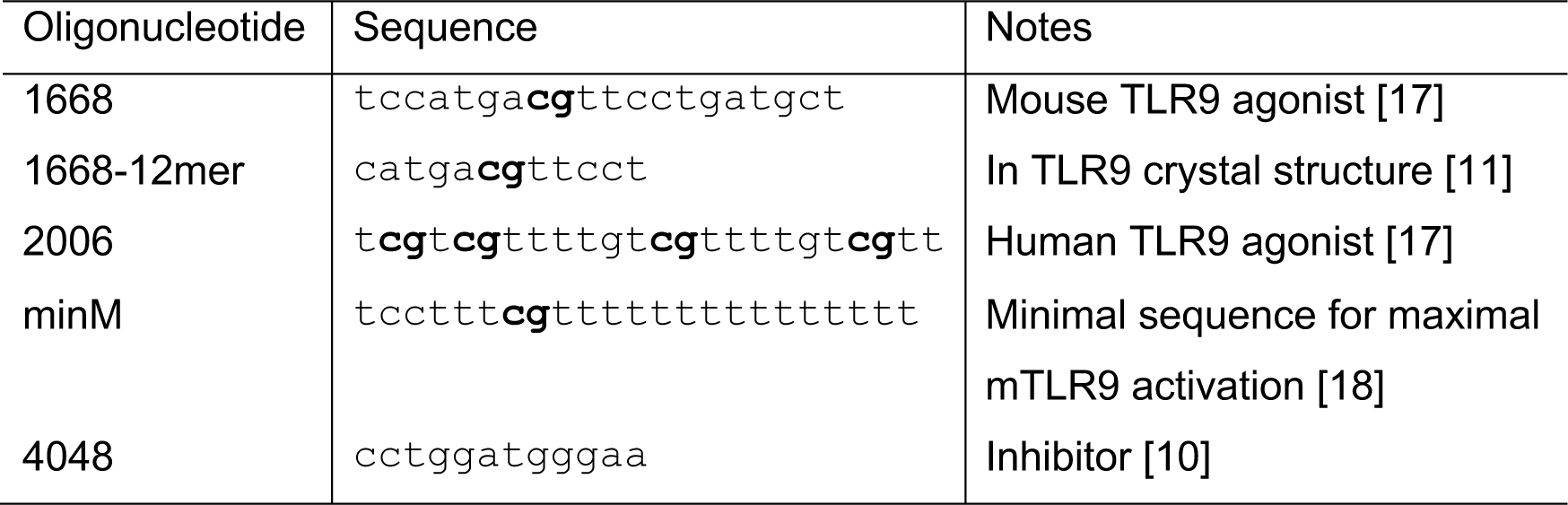
The sequences and properties of TLR9 DNA ssDNA ligands used in this study.

As expected, the hydrodynamic radii and molecular masses calculated from DLS data indicated that mTLR9-ECD formed a 1:1 complex with the antagonist 4084, and 2:2 complexes with all four agonistic oligonucleotides (**Fig. 2A-B**). The experimentally determined molecular diameters of the complexes were slightly larger than expected, and the molecular masses correspondingly smaller, because the DLS data were fitted to a globular model whereas mTLR9-ECD has a non-globular horseshoe shape. The observed range in polydispersity, from 16% to 30%, could be due to small proportions of 1:1 complexes in the TLR9:agonist solutions.

**Fig. 2.**
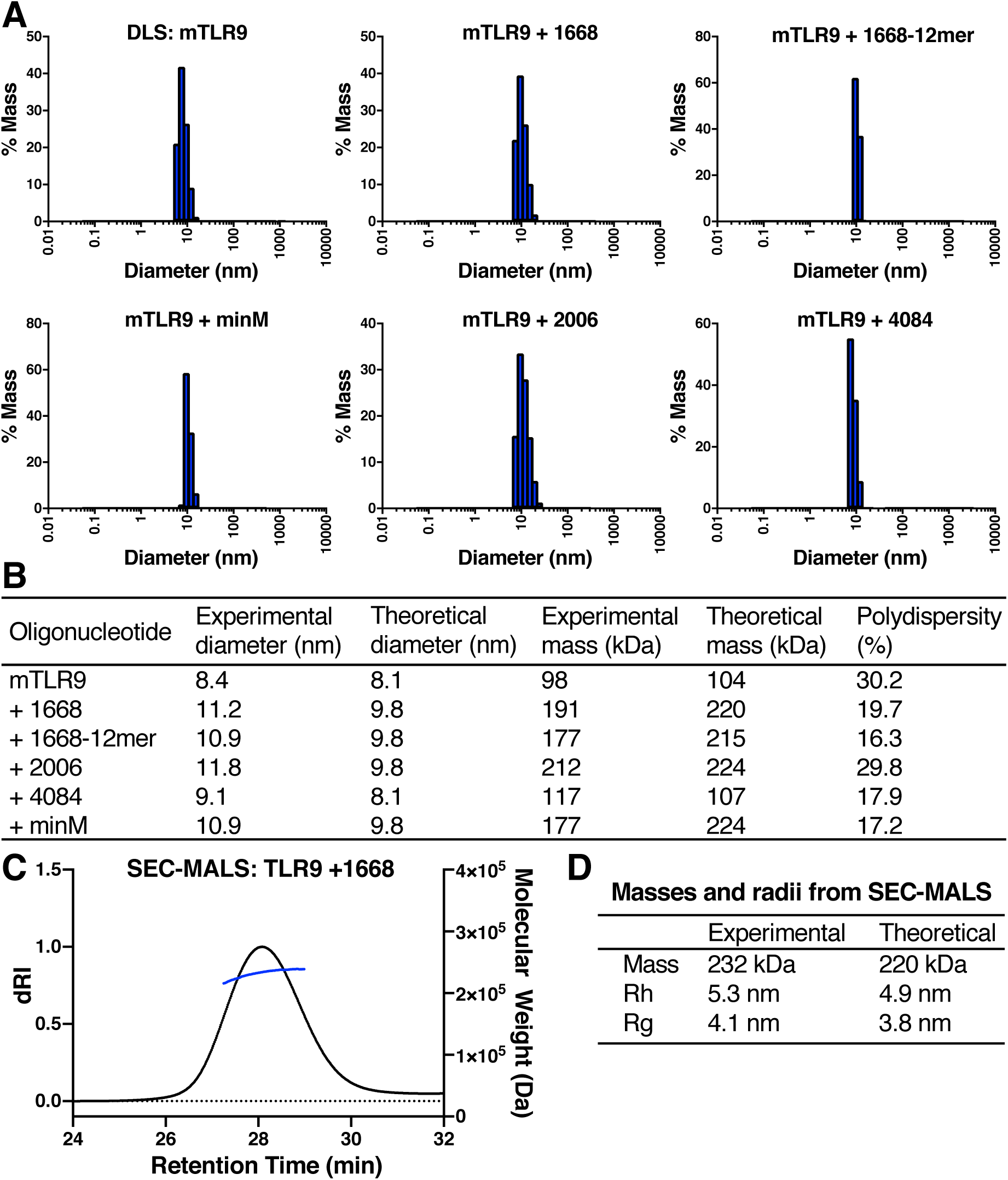
Ligand binding assays with mTLR9 ectodomain (mTLR9-ECD) in the presence of various ssDNA ligands. (**A**) Molecular diameter histograms from dynamic light scattering (DLS). (**B**) Table of experimental and theoretical molecular diameters and masses calculated from DLS data. The polydispersity of each sample, related to the peak width in (A), is listed. The theoretical diameters were calculated as twice radius of gyration, Rg, of monomeric or dimeric TLR9 from the crystal structures [11] divided by 0.775, to convert to diameter of hydration, Rh (assuming Rh = Rg/0.775). (**C**) SEC-MALS of 8 µM mTLR9-ECD with 20 µM oligonucleotide 1668. (**D**) Masses, Rh and Rg determined from SEC-MALS data or calculated from the crystal structure.

To obtain a more direct measure of the mass of a TLR9 dimer, size-exclusion chromatography coupled to multiangle light scattering (SEC-MALS) was performed on mTLR9 bound to oligonucleotide 1668. As expected, the measured mass of 232 kDa was consistent with a 2:2 dimer (**Fig. 2C**). We note that the experimental hydrodynamic radii (Rh) determined from SEC-MALS and DLS (5.3-5.6 nm) were approximately 10% larger than the theoretical radius predicted from the TLR9:1668-12mer crystal structure (4.9 nm; **Fig. 2D**). This slight discrepancy could be due to the method used to calculate Rh (which was based on the root mean square distance from the center of mass), or to the eight additional nucleotides in 1668 versus 1668-12mer, which were not taken into account.

### The equilibrium binding affinity of agonists is not accurately modeled by a 1:1 fit

To further investigate the binding modes of TLR9 ligands, fluorescence polarization (FP) anisotropy ligand binding assays were performed. First, mTLR9-ECD was titrated into 2 nM solutions of oligonucleotides 1668, 1668-12mer, and 4084 labeled with Alexa Fluor 488. The binding curves were fitted with a 1:1 ligand binding model, accounting for receptor depletion (**Fig. 3**). The mTLR9 binding affinities were 28.8 ± 7.2 nM for 4084; 9.2 ± 2.2 nM for 1668-12mer; 5.3 ± 1.4 nM for 1668; and 3.2 ± 0.9 nM for minM. These values are consistent with expectations, since minM is the most potent ligand and 1668-12mer has a shorter than optimal sequence. The FP data fit the 1:1 binding model well for oligonucleotide 4084, which does not induce dimerization. For the agonistic ligands, the data points follow a steeper sigmoidal trajectory than the 1:1 model curve, suggesting that assembly of 2:2 dimer complexes is cooperative. The 2:2 model in **Fig. 1** contains too many variables for fitting to the FP data.

**Fig. 3.**
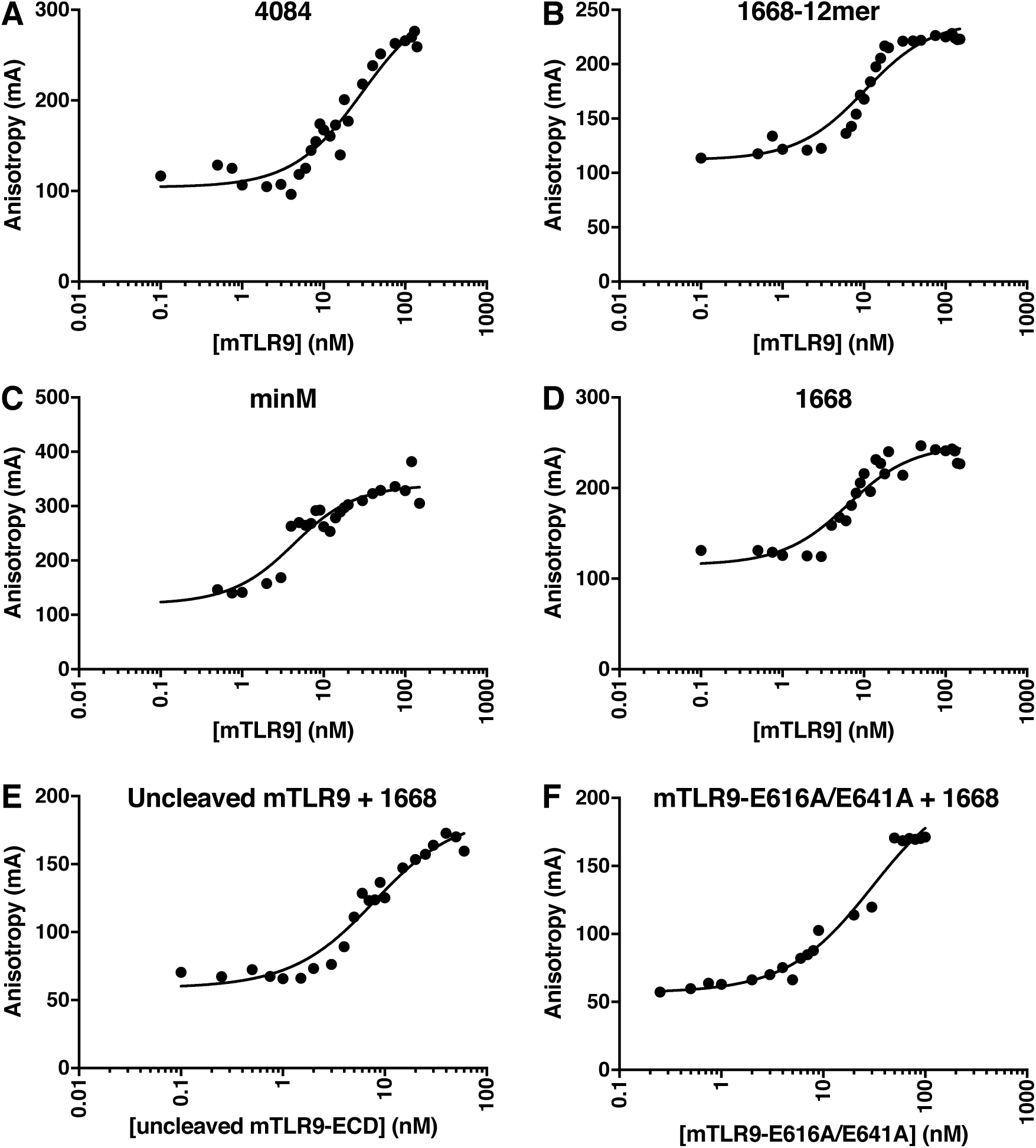
Binding affinities of mTLR9-ECD for various ligands measured by fluorescence polarization (FP). (**A-D**) The equilibrium binding affinities of different Alexa Fluor 488-labeled oligonucleotides for proteolytically activated mTLR9 were calculated from fitting to a 1:1 binding model. (**E**) The affinity of oligonucleotide 1668 for mTLR9-ECD without proteolytic activation. (**F**) The affinity for oligonucleotide 1668 for mTLR9-ECD mutated at Site B.

Cleavage of the ectodomain by an endosomal protease is necessary for dimerization but not for ligand binding [11, 23, 24]. To confirm this in our system, FP was performed with uncleaved mTLR9-ECD and Alexa Fluor 488-labeled 1668. The binding affinity was 6.9 ± 1.5 nM, with a good fit to a 1:1 model curve, consistent with the expected inability of the uncleaved ectodomain to dimerize (**Fig. 3E**). Moreover, since uncleaved TLR9 cannot dimerize, this binding affinity reports on only two of the macroscopic equilibrium constants defined in **Fig. 1**, K_1_ and K_2_. The slight deviation in the data from the theoretical fit (**Fig. 3E**) is likely due to the presence of two ligand binding sites on TLR9, Sites A and B, which the crystal structure suggests have different binding affinities [11]. Hence, early in the titration the ligand will primarily bind the high-affinity site (Site A), with the low-affinity site (Site B) becoming saturated with ligand last.

To deconvolute the contributions of the two ligand binding sites in mTLR9, two key residues involved in ligand binding at Site B were mutated. The mutations, H616A and E641A, are predicted to inhibit ligand binding at Site B. Since the H641A mutation alone abolished TLR9-dependent signaling in a cell-based assay [11], we also predicted that these mutations would inhibit dimerization. Indeed, Alexa Fluor 488-labeled 1668 oligonucleotide bound mTLR9-H616A/E641A with an FP response curve fitting a 1:1 binding model similar to the binding curve for uncleaved mTLR9 (**Fig. 3F**), suggesting that the mutations in Site B prevent dimerization. The binding affinity of mTLR9-H616A/E641A for 1668 was 27.7 ± 6.5 nM. This provides to the affinity of oligonucleotide 1668 for Site A, which corresponds to the microscopic constant K_A_.

### Competition assays suggest a slow dissociation of the dimer

To further characterize the binding mode of TLR9 ligands, competition assays were performed. The binding sites of the 1668-12mer agonist and 4048 antagonist oligonucleotides are partially overlapping [11]. To establish whether these two oligonucleotides bind competitively, a competition experiment was performed by titrating in mTLR9-ECD preincubated with a molar excess of unlabeled 4048 oligonucleotide into a solution containing Alexa Fluor 488-labeled 1668-12mer. No binding of 1668-12mer was observed, indicating that binding of agonist 1668 and antagonist 4084 is competitive (**Fig. 4A**). Next, we examined the equilibrium dynamics of this competition by preincubating mTLR9 with fluorescently-labeled 1668-12mer, titrating in a molar excess of unlabeled 4084, and monitoring displacement of 1668-12mer at various timepoints. Unexpectedly, it took several hours for the competition conditions to reach equilibrium (**Fig. 4B**). This was also true when the same experiment was performed using unlabeled 1668-12mer instead of 4084 as the competing oligonucleotide (**Fig. 4C**). After 4.5 hours, the apparent IC50 of 1668-12mer was approximately 10 nM. Application of the Cheng-Prusoff equation yields an inhibitor constant, Ki, of 8 nM, similar to the Kd for 1668-12mer of 9.2 nM. This agreement between the Kd of labeled ligand and Ki of unlabeled ligand indicates that the fluorescent label does affect binding to mTLR9. Based on the ability of bound oligonucleotides to slowly exchange (**Fig. 4B-C**), we conclude that oligonucleotides dissociate from the dimer very slowly, on the timescale of hours.

**Fig. 4.**
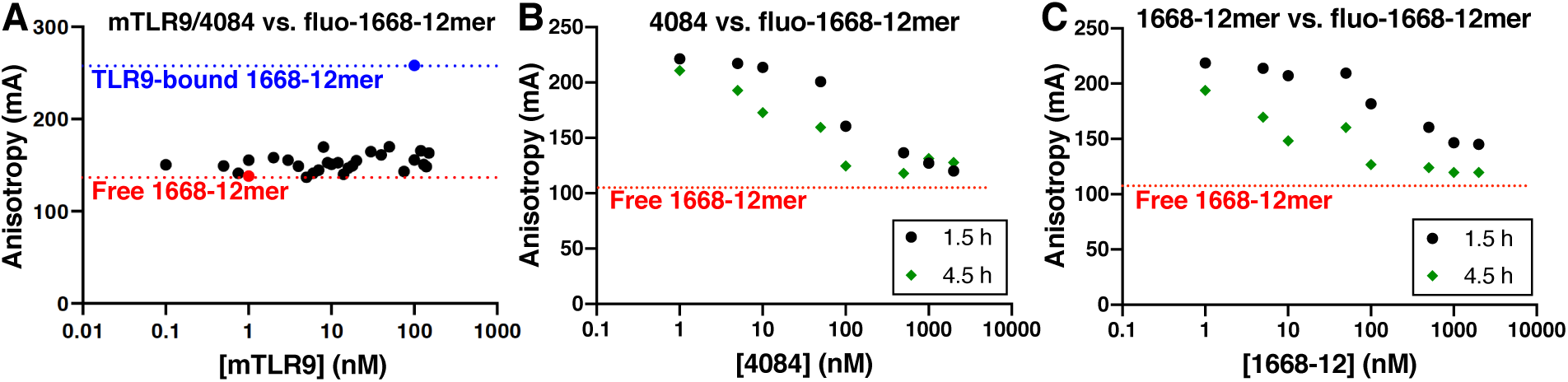
Competition FP experiments reveal a slow dissociation of TLR9 dimers. (**A**) mTLR9-ECD preincubated with 2 µM unlabeled oligonucleotide 4048 was titrated into a solution containing 5 mM Alexa Fluor 488-labeled 1668-12mer oligonucleotide. The blue dotted line marks the anisotropy of 1668-12mer bound to mTLR9 in the absence of 4048. The red dotted line marks the anisotropy of free fluorescent 1668-12mer. (**B-C**) Unlabeled 4084, (**B**), or 1668-12mer, (**C**), was titrated into a solution containing 100 nM mTLR9-ECD preincubated with 2 nM fluorescent 1668-12mer. The red dotted line marks the anisotropy of the fluorescent 1668-12mer, which is the expected anisotropy if all of the fluorescent oligonucleotide has been competed off the protein. Measurements were taken after 1.5 h (black) and 4.5 h (green).

## Discussion

Here we present a robust theoretical equilibrium binding model for TLR9 binding to DNA ligands complemented by *in vitro* biophysical data on the TLR9 ectodomain binding to ligands. Our DLS and SEC-MALS ligand binding assay confirm that agonistic oligonucleotides induce dimerization of recombinant, proteolytically activated mTLR9-ECD, whereas TLR9 remained monomeric in the presence of antagonistic ligand 4084. Fluorescence polarization anisotropy experiments showed that all ligands bound TLR9 tightly with overall apparent Kd values in the low nanomolar range. More importantly, binding of ligands that induced TLR9 dimerization did not fit a 1:1 binding model, consistent with a more complex binding mode. Moreover, the unexpectedly long time that it took ligand competition experiments to reach equilibrium (several hours) revealed that dissociation of oligonucleotides from dimers is very slow, despite the rate of dimer assembly being relatively rapid. We speculate that oligonucleotide dissociation would be accompanied by TLR9 dimer dissociation.

Binding assays with a TLR9 variant containing mutations at Site B provided clear evidence that both oligonucleotides binding sites are required for dimerization and provided the microscopic constant for ligand binding to Site A, K_A_ (28 nM). Together, these experiments and the DLS data for apo TLR9 provide experimental evidence that K_3_ > K_1_. Performing the analogous experiment with an mTLR9 variant mutated at Site A would provide the microscopic binding constant for ligand binding to Site B, K_B_. With both K_A_ and K_B_ known, ligand binding curves for uncleaved TLR9, an obligate monomer, could be fitted to our 2:2 binding model to determine whether Sites A and B are in fact independent and non-cooperative as suggested by the crystal structure.

Our ligand binding studies were performed in the absence of auxiliary oligonucleotides (5’-xCx), which were recently shown to augment signaling. The purpose of this study was to develop an accurate model for TLR9 binding to agonistic oligonucleotides and including auxiliary oligonucleotides would have complicated interpretation of ligand binding data. However, the role of auxiliary oligonucleotides is an important area for further study. In particular, it will be important to examine whether 5’-xCx oligonucleotide binding at the auxiliary site is independent of ligand binding at Sites A and B, and to determine the mechanism through which auxiliary ligands promote dimerization.

In summary, a complete model for TLR9 binding is presented, and while there are many solutions for the macroscopic equilibrium constants *a priori*, the experimental data presented narrows the relationships between the macroscopic binding constants. To obtain a unique solution for the complex 2:2 binding model of TLR9 to its ligands, further experimental or numerical analyses are required. Since the activation of TLR9 is similar to the activation of other TLRs, most notably TLR7 and TLR8, this work will help establish a more general model of TLR activation, and hopefully guide future efforts to design TLR9 agonists or antagonists.

## Materials and Methods

### Expression and purification of recombinant mouse TLR9 ectodomain (mTLR9-ECD)

A pMT plasmid encoding mTLR9-ECD with a secretion signal and a C-terminal protein A tag was a kind gift from Prof. Toshiyuki Shimizu. The Site B mutant was generated by site-directed mutagenesis. The plasmid was co-transfected into S2 insect cells with 10:1 molar excess of pCoBlast using TransIT-Insect transfection reagent (Mirus Bio). Stable cell lines were selected with 100 µg/mL blasticidin (Cambridge Bioscience). Protein expression was induced with 0.5 mM copper sulfate. Five days after induction, the media containing secreted mTLR9-ECD was concentrated by tangential-flow filtration on a 30 kDa cut-off membrane (Merck). mTLR9-ECD was purified by protein A-affinity chromatography with IgG Sepharose 6 Fast Flow resin (GE Healthcare), washed with PBS and eluted with 0.1 M glycine-HCl pH 3.5, 0.15 M NaCl, and immediately neutralized with 1/20 (v/v) 1 M Tris pH 8. mTLR9-ECD was further purified by ion-exchange chromatography with a monoS 4.6/100 PE column (GE Healthcare) in 10 mM MES pH 6.0, 0.06 – 1 M NaCl. Protein eluting at 0.25-0.32 M NaCl was pooled, cleaved, and further purified by size-exclusion chromatography (SEC) on a Superdex 200 10/300 column (GE Healthcare) in 10 mM Tris pH 7.4, 0.15 M NaCl. Uncleaved protein eluted as a mixture of monomer and dimer in SEC. To remove the tag and proteolytically activate mTLR9-ECD, 1/20-1/50 (w/w) GluC protease (New England BioLabs) was added and incubated 24-48 h at 4°C. GluC was removed with benzamidine Sepharose 4 Fast Flow (GE Healthcare) resin. The cleaved mTLR9-ECD eluted as a monomer in SEC.

### Dynamic light scattering (DLS)

2 µM mTLR9-ECD was mixed with 2 µM ssDNA oligo (Sigma-Aldrich) in 10 mM MES pH 6, 0.15 M NaCl. Following a one hour, room temperature incubation, the samples were filtered through a 0.22 µm spin filter (Costar) and 30 µL were loaded into a black, clear bottom, 384 well plate (Corning). Dynamic light scattering was collected on a Wyatt Technologies DynaPro II plate reader at 25°C. 5 acquisitions were collected for each sample, with five measurements per acquisition.

### Size-exclusion chromatography and multiangle light scattering (SEC-MALS)

8 µM mTLR9-ECD was incubated with 20 µM oligonucleotide 1668. 0.1 ml of this solution was subjected to SEC at 293 K on a Superdex 200 10/300 column (GE Healthcare) preequilibrated in 10 mM MES pH 6.0, 0.15 M NaCl with a flow rate of 0.5 ml min-1. Protein in the eluate was detected with a UV detector at 280 nm (Agilent Technology 1260 UV), a quasi-elastic light scattering (QELS) module (DAWN-8+, Wyatt Technology) and a differential refractometer (Optilab T-rEX, Wyatt Technology). Molar masses of peaks in the elution profile were calculated from the light scattering and protein concentration, quantified using the differential refractive index of the peak assuming dn/dc = 0.186, using ASTRA6 (Wyatt Technology).

### Fluorescence polarization anisotropy assays

For ligand binding assays, recombinant mTLR9-ECD was titrated into a solution of oligonucleotide labeled at the 5’ end with Alexa 488 Fluor (Sigma-Aldrich). The oligonucleotide concentration, either 2 nM or 5 nM, was optimized to ensure that the observed increase in fluorescence was due to binding and not ligand saturation. A blank was measured in the absence of fluorescent oligonucleotide and with the highest concentration of protein (100 nM). 30 µL samples were assayed in 384-well black, clear-bottomed plates (Corning) with a ClarioSTAR plate reader (BMG Labtech) using the 482/530 nm filter. The gain and focal point were adjusted to ensure the readings were within an appropriate range.

Data were fitted with a 1:1 model for binding that accounted for ligand depletion [25], using the following equation:

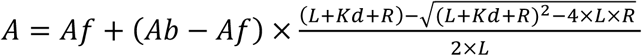

Where *Af* denotes the anisotropy of the free ligand, *Ab* the anisotropy of the bound ligand, *L* the total concentration of ligand and *R* the total concentration of protein. The concentration of ligand (*L*) was fixed, and the other parameters, *Ab, Af*, and *Kd* were fitted using the known values of *R* and *A*. The fit was performed with Prism8 (GraphPad).

For competition assays where an unlabeled oligonucleotide was displacing a bound fluorescent oligonucleotide, 100 nM mTLR9-ECD and 2 nM fluorescent oligonucleotide were preincubated for 30 min at room temperature in 10 mM MES pH 6, 0.15 M NaCl. Unlabeled competing oligonucleotide was titrated in. Measurements were taken at 1.5 h and 4.5 h. For competition assays where a fluorescent oligonucleotide was displacing an unlabeled oligonucleotide, 2 µM oligonucleotide 4084 at was preincubated with increasing amounts of mTLR9-ECD. Following a brief incubation at room temperature, 2 nM or 5 nM of the fluorescent oligonucleotide was added. The inhibitor constant, *Ki*, was calculated with the Cheng-Prusoff equation:

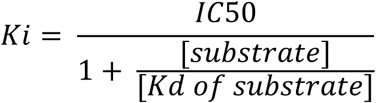

## Abbreviations

CpG: unmethylated CG DNA nucleotide sequence motif;
DLS: dynamic light scattering;
LRR: leucine-rich repeat;
mTLR9-ECD: mouse Toll-like receptor 9 ectodomain;
PAMP: pathogen-associated molecular pattern;
sDNA: single-stranded DNA;
SEC-MALS: size-exclusion chromatography coupled to multiangle light scattering;
TIR: Toll/interleukin18 receptor domain.

## Author contributions

Conceptualization, S.R. and Y.M.; Methodology, all authors; Investigation, S.R., C.M.J. and S.H.M.; Writing – Original Draft, S.R. and Y.M.; Writing – Review & Editing, all authors; Visualization, S.R. and Y.M.; Supervision, Y.M.; Project Administration, Y.M.; Funding Acquisition, Y.M.

## Acknowledgements

We thank Prof. Toshiyuki Shimizu (University of Tokyo) for kindly providing TLR9 cDNAs. We thank members of the Modis lab for insightful discussions. This work was supported by NIH grant R01 GM102869 and Wellcome Trust Senior Research Fellowship 101908/Z/13/Z to Y.M.

